# EEG-based personal identification: comparison of different functional connectivity metrics

**DOI:** 10.1101/254557

**Authors:** Matteo Fraschini, Luca Didaci, Gian Luca Marcialis

## Abstract

Growing interest is devoted to understanding how brain signals recorded from scalp electroencephalography (EEG) may represent unique fingerprints of individual neural activity. In this context, the present paper aims to investigate the impact of some of the most commonly used techniques to estimate functional connectivity on the ability to unveil personal distinctive patterns of inter-regional interactions. Different metrics, commonly used to estimate functional connectivity and derived centrality measures, were compared in terms of equal error rate. It is widely accepted that each metric carries specific information in respect to the underlying interactions network. Nevertheless, the reason why these metrics convey different subject specific information has not been investigated yet. Experimental results on two publicly available datasets suggest that different functional connectivity metrics define a peculiar subjective profile of connectivity and have different mechanisms to detect subject-specific patterns of inter-channel interactions. It is important to consider the effects that frequency content and spurious connectivity values may play in determining subject-specific characteristics.

## 1. Introduction

The interest towards the investigation of subject specific human characteristics that can be used to develop robust biometric systems still represents a big challenge. In this context, growing interest is devoted to understanding how brain signals recorded from scalp electroencephalography (EEG) may represent a unique fingerprint of individual neural activity. In the last few years a huge number of works have investigated the potential role of EEG signal characteristics as biometric system (about 300 new papers in the last 10 years). A detailed literature overview of the proposed methods is therefore quite challenging and in any case out of the scope of the present paper. Nevertheless, some attempts to summarize the state of the art was previously proposed in (Campisi and La Rocca, 2014; Khalifa et al., 2012; Del Pozo-Banos et al., 2014). In brief, it is possible to consider the approaches proposed so far mainly organized into two fundamental categories: (i) task based and (ii) resting-state based EEG analysis. The first category is oriented on experimental setups that allow to investigate properties of the EEG signal that are strictly related to the onset of a specific stimulus. Motor (real and imagery) tasks (Yang et al., 2018), visual evoked potentials (Das et al., 2015; Palaniappan and Mandic, 2007; Armstrong et al., 2015), auditory stimuli (Light et al., 2010), imagined speech (Brigham and Kumar, 2010), eye blinking (Abo-Zahhad et al., 2016) and multiple functional brain systems (Ruiz-Blondet et al., 2016) have been proposed so far in order to elicit individual unique responses. In contrast, the second category is mainly oriented to detect characteristic patterns of induced brain activity at rest (both during eyes-closed and eyes-open. In line with the extensive use of tools from modern network science to understand brain complex organization (Stam, 2014), measure of functional connectivity (La Rocca et al., 2014; DelPozo-Banos et al., 2015; Han et al., 2015; Garau et al., 2016) and network metrics have been recently proposed (Crobe et al., 2016; Fraschini et al., 2015) as EEG-based biometric traits. However, it seems still evident that there exists a gap between current investigations of EEG signal as neurophysiological marker and its application in personal verification systems. In particular, it is widely accepted that each metric used to assess functional connectivity carries specific information with respect to the underlying interactions network (Kida et al., 2016). Nevertheless, the reason why these metrics convey different subject specific information has not been investigated yet. Following what reported in (Garau et al., 2016; Fraschini et al., 2015), the present paper aims to investigate and compare the impact of some of the most commonly used techniques to estimate functional connectivity on the ability to detect personal unique discriminative features based on inter-regional interaction profiles. In order to answer this question, we focused our attention on measures based on different properties of the original signal. In particular, the following measures were included in the present study: (i) the Correlation Coefficient (CC); (ii) the Phase Lag Index (PLI) (Stam et al., 2007), which quantifies the asymmetry of the distribution of phase differences between two signals; (iii) the corrected (after performing the orthogonalisation of raw signals) and (iv) the uncorrected version of Amplitude Envelope Correlation (AEC) (Hipp et al., 2012; Brookes et al., 2011), which provides functional coupling without coherence or phase coherence; (v) the corrected and (vi) the uncorrected version of the Phase Locking Value (PLV) (Lachaux et al., 1999), which detects frequency-specific transient phase locking independently from amplitude. Each of the proposed metrics has different properties and capture different characteristics of the EEG signal interactions which will be discussed in this paper. We hypothesized that the choice of the metric may have a great impact in highlighting subject specific patterns of functional interactions, and that advantages and disadvantages of each technique should be correctly taken into account when interpreting the corresponding results in terms of performance of a biometric verification system. Finally, the comparison was also performed at the level of nodal relevance, as expressed using two different network centrality measures, namely node strength and eigenvector centrality.

## 2. Material and methods

### 2.1 EEG datasets

The analysis was performed using two different EEG datasets. The first one (DS01) is a freely available EEG dataset containing 64 channels scalp recordings from 109 subjects with eyes-closed and eyes-open resting-state conditions, each lasting 1 minute of spontaneous brain activity. The full dataset was created and contributed to PhysioNet (Goldberger et al., 2000) by the developers of the BCI2000 instrumentation system. A detailed description of the original system can be found in (Schalk et al., 2004) and access to the raw EEG recording is possible at the following website: https://www.physionet.org/pn4/eegmmidb/. For the purpose of the present paper our analysis was applied to resting state eyes-closed condition. The second dataset (DS02), including eyes-closed anonymized EEG from 11 subjects, is freely available without restriction and was released of the authors of a previous study (Sockeel et al., 2016). The EEG acquisitions were carried out with a 62-channel BrainAmp cap with sampling frequency of 5 kHz. The reference electrode was located on Cz channel, the ground electrode below Oz and the electrode impedance did not exceed 10kOhm. As for the original work, only the last 300 seconds were considered for the subsequent analysis.

### 2.2 Preprocessing

For both the datasets, original raw data were band-pass filtered in the common frequency bands: delta (1 - 4 Hz), theta (4 - 8 Hz), alpha (8 - 13 Hz), low-beta (13 - 20 Hz), high-beta (20 - 30Hz) and gamma (30 - 45 Hz). Successively, each single EEG recording was organized into five different epochs (without overlap) of 12 seconds (Fraschini et al., 2016).

## 3. Connectivity metrics

From the preprocessed EEG signals, separately for each subject, each epoch and each frequency band, a connectivity matrix was computed. Each single entry of the connectivity matrix, which represents the weight of the functional interaction, was evaluated by using the following different metrics.

### 3.1 Correlation

The Correlation Coefficient (CC) represents the simpler method to estimate statistical relationship between two random variables and it is widely used in fMRI studies (Friston et al., 1994). However, since scalp EEG signals contain electric fields derived from common current sources, CC does not represent the optimal metric to estimate functional interactions in this context. In this study, CC was mainly applied in order to quantify the possible effect of spurious patterns of connectivity on the definition of subject specific EEG traits.

### 3.2 Phase lag index

The phase lag index (PLI) (Stam, 2014) is a technique that quantify the asymmetry of the distribution of phase differences between two signals and removes the effect of amplitude information. Furthermore, PLI is less affected by the influence of common sources and thus defines more reliable interactions between the underlying signals. The PLI is computed as the asymmetry of the distribution of instantaneous phase differences between two signals:

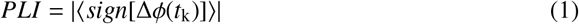

### 3.3 Amplitude Envelope Correlation

Band limited amplitude envelop correlation (AEC) (Hipp et al., 2012; Brookes et al., 2011), using Hilbert transform, was also used in this study. In particular, the envelope is obtained by measuring the magnitude of the analytic signal and successively the Pearson’s correlation between envelopes is computed as a metric of functional connectivity.

### 3.4 Amplitude Envelope Correlation, corrected version

It is well known that signal components that pick up the same source at different sites (i.e., EEG channels) have an identical phase. In this work, to overcome this possible limitation, we used an orthogonalisation procedure performed in the spatial domain (by removing the linear regression) before to compute the AEC values. In the present paper, the corrected version of AEC is reported as AECc.

### 3.5 Phase Locking Value

The phase locking value (PLV), introduced by (Lachaux et al., 1999), allows to detect transient phase locking values which are independent of the signal amplitude. The PLV therefore represents the absolute value of the mean phase difference between the two signals:

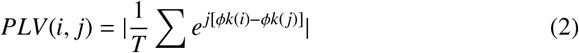

### 3.6 Phase Locking Value, corrected version

As for the AEC, PLV may still provide spurious zero-lag connectivity between different EEG signals, therefore the same orthogonalisation procedure was also applied to PLV values. In the present paper, the corrected version of PLV is reported as PLVc.

### 3.7 Node centrality

In order to evaluate the node relevance within each network, two different centrality measures were computed from each connectivity matrix. The node strength (S), which represents the sum of weights of links connected to the node, and the eigenvector centrality (EC), which is a self-referential measure of centrality where nodes have higher centrality if they are connected to nodes that have high centrality.

## 4. Performance evaluation

The performance obtained by the application of the different connectivity metrics for each of the two datasets have been reported in terms of Equal Error Rate (EER). The EER refers to the rate at which both acceptance error (that occurs when the system accepts an impostor) and rejection error (that occurs when the system rejects a genuine match) are equal. It represents a quick way to compare the accuracy of different systems and it is widely used in evaluating the performance of biometric fingerprints. In short, the EER is the point where false identification and false rejection rates are equal, thus the lower the EER, the better the performance of the system. As previously proposed (Fraschini et al., 2015), the system performance is based on the computation of genuine and impostor matching scores. The scores, computed separately for each frequency band, represent the Euclidean distance (d) between each pair of feature vectors. The feature vectors are represented by the connectivity values extracted from the upper triangular symmetrical connectivity matrix obtained by using the different metrics. Thus, each feature vector contains (number of channels) x (number of channels - 1) / 2 elements, each representing the corresponding statistical interdependence between pairs of EEG channels. Finally, from the matching scores, the similarity scores have been computed as 1 / (1 + d), where d represents the Euclidean distance.

## 5. Results

The Results section is organized into three main sub-sections. In the first sub-section (A), we reported the results obtained using the first dataset (DS01), the second sub-section (B) contains the results from the replication analysis based on the second dataset (DS02) and the third sub-section (C) deals with the analysis on the centrality measures extracted from the FC matrices.

### 5.1 Dataset: DS01

The results obtained on the first EEG dataset (DS01) are depicted in Figure 1 (which represents for each connectivity metric the corresponding average matrix and standard deviation) and summarized Table 1. These findings highlight a tendency to obtain higher performance (lower EER) for the higher frequency bands. This result confirms previous findings (Garau et al., 2016; Fraschini et al., 2015) which suggested a potential role of myogenic activity, which is known to affect high frequencies (Muthukumaraswamy, 2013), on the definition of individual EEG characteristics. Moreover, these reported results do not show any potential effect induced by the use of metrics which correct for signal spread.

**Figure 1.**
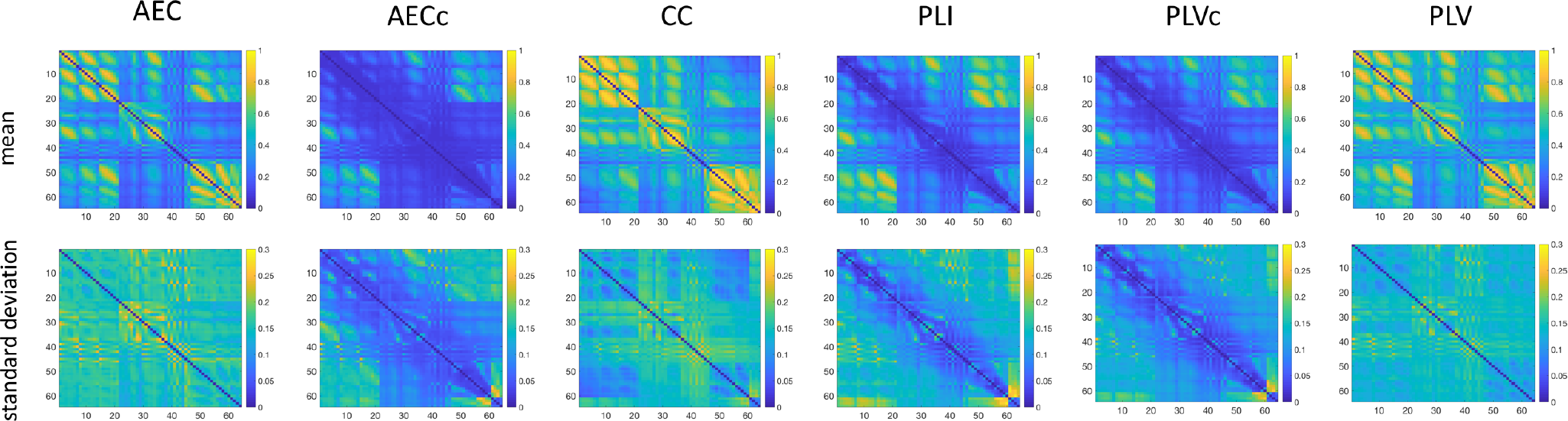
Connectivity patterns for each metric and corresponding between subjects variation expressed as standard deviation.

**Table 1.** The performance of different connectivity metrics expressed as EER for each frequency band obtained using the first EEG dataset (DS01). EER values lower than 10% were marked as bold.

|  | AEC | AECc | CC | PLI | PLVc | PLV |
| --- | --- | --- | --- | --- | --- | --- |
| Delta | 46.37 | 24.12 | 17.14 | 44.15 | 26.14 | 42.25 |
| Theta | 43.43 | 25.36 | 17.42 | 41.06 | 31.83 | 38.62 |
| Alpha | 40.30 | 23.40 | 15.10 | 31.22 | 26.69 | 31.49 |
| Low Beta | 25.82 | 11.66 | **9.04** | 16.61 | 21.15 | 15.21 |
| High Beta | 16.87 | **8.18** | **6.63** | **8.47** | 16.80 | **8.31** |
| Gamma | 13.86 | **9.77** | **6.88** | **5.70** | 11.02 | **5.91** |

### 5.2 Dataset: DS02

The results obtained on the second EEG dataset (DS02) are summarized Table 2. These findings tend to confirm the results obtained on the first EEG dataset (DS01). In particular, this replication analysis still shows a strong effect of the frequency content on the system performance, as the better results (lower EER) are obtained for high frequency bands (high-beta and gamma). Furthermore, these findings suggest that the FC metrics robust to the effect of linear mixing/signal spread, in particular AECc and PLVc, show worse performance even for high frequency bands.

**Table 2.** The performance of different connectivity metrics expressed as EER for each frequency band obtained using the first EEG dataset (DS02). EER values lower than 10% were marked as bold.

|  | AEC | AECc | CC | PLI | PLVc | PLV |
| --- | --- | --- | --- | --- | --- | --- |
| Delta | 42.31 | 49.80 | 33.07 | 47.04 | 48.16 | 28.76 |
| Theta | 33.75 | 45.56 | 30.38 | 47.91 | 49.13 | 33.11 |
| Alpha | 29.96 | 43.58 | 20.16 | 41.53 | 40.62 | 17.49 |
| Low Beta | 26.07 | 44.11 | 23.42 | 42.16 | 42.15 | 11.69 |
| High Beta | 22.18 | 45.20 | 12.91 | 42.35 | 42.40 | **6.62** |
| Gamma | 17.31 | 36.78 | 14.67 | 11.65 | 16.53 | **3.07** |

### 5.3 Centrality

The results obtained from the centrality measures as assessed by using the node strength (S) and eigenvector centrality (EC) are shown in Table 3 for both DS01 and DS02 dataset. The results refer only to the gamma band since this was the frequency content that shows the best results in terms of EER over all the other frequencies.

**Table 3.**
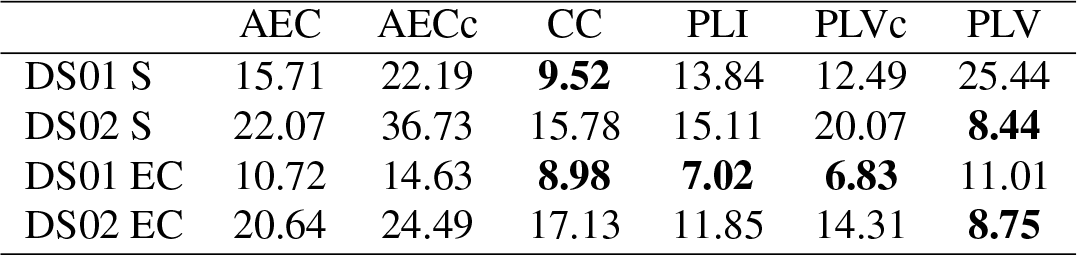
The performance of different centrality measure for each connectivity metric expressed as EER for the gamma frequency band. S refers to strength and EC to eigenvector centrality. EER values lower than 10% were marked as bold.

## 6. Discussion

The present paper aimed to investigate and compare the impact of common techniques used to estimate functional connectivity on their capacity to detect personal discriminative features. In summary, the results of the present paper show, as expected, that different metrics, each characterized by different mechanisms of functional interaction, define a peculiar subjective profile of connectivity. In our opinion, two main points deserve particular attention. The first important point is related to a marked association between the frequency content and the ability to discriminate among different subjects (as reported using the EER). Indeed, for both the EEG datasets the best performance (lower EER) were obtained for the higher frequency bands (high beta and gamma). It is interesting to note that this finding represents a confirmation of previously reported results using different approaches (Crobe et al., 2016; Fraschini et al., 2015). In this context, it is not possible to rule out the hypothesis that muscle artifacts, particularly evident at high frequencies (Muthukumaraswamy, 2013), may play a key role in the definition of discriminative (individual) characteristics. The second point is related to the different performance obtained using the different connectivity estimators. It is evident that some of connectivity metrics, namely AEC (corrected and uncorrected versions) and the corrected PLV, give the worst performance even for the higher frequency bands. This phenomenon is still more clear from the results obtained using the second dataset (DS02), where the corrected versions of AEC and PLV do not exceed respectively 35% and 16% of EER, where their counterparts (not corrected versions) show much lower EER, with values respectively up to 17% and 3%. Furthermore, this latest result represents the best performance obtained over the entire analysis. These findings are also confirmed when the centrality measures were extracted from the connectivity metrics. Again, the worst results were obtained for AEC metrics. In general, the second dataset (DS02) shows worst results if compared with the first dataset (DS01). However, it is important to note that, as it is evident from visual inspection of EEG traces, the second dataset is certainly less affected by EEG artifacts. This aspect, in our opinion, strengthen the reported results since the first dataset (which is particularly noisy) better represents real-life EEG acquisitions, thus giving a better simulation of possible real biometric applications.

Moreover, considering the inherently different characteristics of the used EEG datasets, this study shows that some connectivity metrics, namely non-corrected version of PLV, are more prone to pick up distinctive EEG features even when the original signals are particularly free from artifacts and contaminations. The other way around, it should be noted that PLV is a connectivity metrics that is deeply influenced by mechanisms of volume conduction, signal spread and common sources. Therefore, caution should be used when interpreting the reported results. In particular, it is still possible that the distinctive patterns of connectivity, as highlighted by PLV, may be strongly influenced by spurious connectivity values generated by the previously cited sources of noise (i.e., volume conduction, signal spread and common sources).

## 7. Conclusions

Future works should investigate if the results reported so far at scalp level still hold when the EEG signals are reconstructed (by resolving the inverse problem) at source level where the effects due to spurious connections, at least in part, attenuated. Finally, this work suggests that different functional connectivity metrics have different mechanism to detect subject specific patterns of inter-channel interactions, that it is important to consider the effect of the frequency content and that spurious connectivity values may play an important role in this context.

